# Which perceptual categories do observers experience during multistable perception?

**DOI:** 10.1101/2024.09.24.614648

**Authors:** Jan Skerswetat, Peter J. Bex

## Abstract

Multistable perceptual phenomena provide insights into the mind’s dynamic states within a stable external environment and the neural underpinnings of these consciousness changes are often studied with binocular rivalry. Conventional methods to study binocular rivalry suffer from biases and assumptions that limit their ability to describe the continuous nature of this perceptual transitions and to discover what kind of percept was perceived across time. In this study, we propose a novel way to avoid those shortcomings by combining a continuous psychophysical method that estimates introspection during binocular rivalry with machine learning clustering and transition probability analysis. This combination of techniques reveals individual variability and complexity of perceptual experience in 28 normally sighted participants. Also, the analysis of transition probabilities between perceptual categories, i.e., exclusive and different kinds of mixed percepts, suggest that interocular perceptual competition, triggered by low-level stimuli, involves conflict between monocular and binocular neural processing sites rather than mutual inhibition of monocular sites.

**Layman abstract:** When our brain receives ambiguous information about the world, it changes its interpretation between different alternatives and thereby provides insight into how the mind works. Scientists often use a technique called binocular rivalry, where each eye sees a different image, to provoke an ambiguous visual world that is perceived as ongoing competition among interpretations of the two eyes inputs. Traditional methods for studying binocular rivalry struggle to describe the continuous nature of this fluctuation and to estimate the range of different perceived experiences. We have created a new approach in which participants reproduce their ongoing perceptual experiences combined machine learning analyses of these states. We found that individuals visual experience is more varied and complex than previously thought. Our results suggest that when our eyes see conflicting images, the brain’s effort to make sense of what is seen involves syntheses among both monocular and binocular brain areas, not just competition between monocular areas.

## Introduction

The quest to understand consciousness has seen a boom of visual paradigms to investigate the relationship between awareness and neural correlates. Methods that provoke endogenous multistable perceptual competition without exogenous stimulus change have become prominent tools to investigate changes of the contents of visual consciousness over time in the minds of humans (1) and other primates. (2) In one multistability paradigm, when different images are presented to each eye viewers typically experience *binocular rivalry* and perceive transitions among the image presented to the left eye (left exclusivity), the image presented to the right eye (right exclusivity) and mixtures of those images (including superimposition and piecemeal combinations). The measurement of binocular rivalry has the potential to identify clinical biomarkers of neuro-atypicality (3) and personality traits. (4)

A well-known problem with the study of these correlates of conscious experience is that the gradual nature of perceptual changes is not well-captured with standard paradigms that are used to measure multistability. During conventional alternative-forced-choice (AFC) tasks, the observer is instructed to classify moment-to-moment changes in their subjective experiences typically by pressing buttons assigned to different perceptual categories. The available categories are pre-selected by the experimenter, are often only described verbally, and have included two exclusive percepts (2 AFC), (5) two exclusive percepts and all mixed percepts (3 AFC, see Figure 1A), (6) two exclusive and two mixed (piecemeal and superimposed) percepts (4 AFC), (7) two exclusive and three mixed (left-predominant, right-predominant or equal superimposition) percepts (5 AFC). (8) The instructions for the perceptual categorization given by the experimenter may further vary between ‘predominance’ (9) and ‘exclusivity’ (10) within each category and even lead to additional judgement criterion of proportions within any moment of viewing (e.g. ≥ 75% predominance (11)). These methods do not provide validated, personalized estimation of perceptual states or their boundaries, make assumption that the experiences described by the experimenter represent the experiences for the participant, do not capture all mixed perceptual experiences reported in the literature, button press methods provide are low in data resolution, nor are they able to track perceptual experiences within mixed categories (piecemeal and superimposition). For further review on rivalry methods please see. (12)

**Figure 1:**
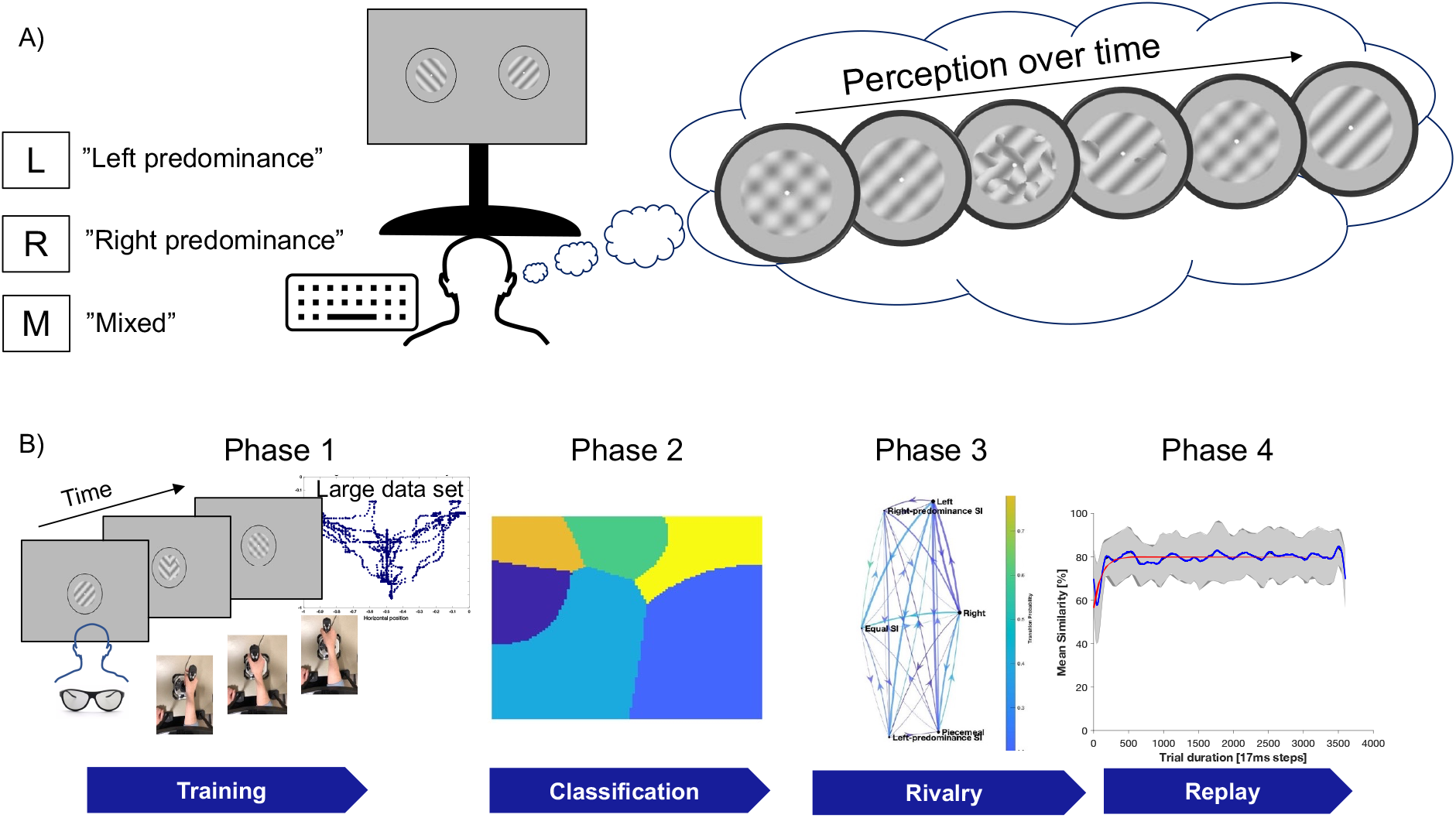
Scheme for different binocular rivalry paradigms. A) Scheme of typical binocular rivalry setting. When an observer views dissimilar stimuli, e.g., achromatic sinusoidal gratings tilted -45° and +45° viewed dichoptically, perceptual competition arises in which experiences gradually change across time, known as binocular rivalry. The traditional task for the observer is then to continuously report what is seen via key presses assigned to categories by an experimenter, here a 3-Alternative-Forced-Choice task. B) Schematic overview of the InFoRM: Rivalry paradigm. During Indicate-Me (Phase 1), participants explore the stimulus-space, moving a joystick to modify binocular-non-rivaling stimuli in real-time that generate corresponding changes of the physical image. The participants were then asked to move the joystick to highlight images that they consider representative of six canonical rivalry states (‘exclusive left-tilted’, ‘exclusive right-tilted’, ‘piecemeal’, ‘equal superimposition’, ‘superimposition with left-tilted predominance’, and ‘superimposition with right-tilted predominance’), that have been reported in previous rivalry literature. During Follow-Me (Phase 2), participants moved the joystick to match perceptual reports for physically changing binocular-non-rivaling-stimuli to confirm their understanding of the relationship between the joystick position and stimulus appearance. Participants followed four trials that reproduced the rivalry experiences of author JS and four trials that reproduced the six rivalry states the participant had generated themselves during phase 1 - Indicate-Me. This trained participants to track their changing experiences during perceptual rivalry while also capturing the participant’s joystick position for each of six canonical perceptual categories. These data used to build estimates of introspection for each category, indicated with different colors in the classification figure. During Rival-Me (Phase 3), participants reported their perception with the same instruction as for Phase 2. The resulting data were then analyzed with various techniques, including the illustrated Hidden Markov Models. During Replay-Me (Phase 4), participants’ responses during the Phase 3-Rival-Me dichoptic-trials were used to generate physically changing binocular stimuli, that the participant again tracked which validated their individual perceptual-state-space. These data from Phase 3 and Phase 4 were then analyzed for similarity illustrated by the plot for one representative participant.

Concerns that the active report requirement of AFC paradigms may unintentionally influence conscious experience have been addressed by no-report paradigms. In these approaches, an observer performs a binocular rivalry task twice: once with and once without an AFC button pressing task while pupil diameter, (13) optokinetic nystagmus, (14) or active gaze changes are simultaneously recorded. (15) The ocular biomarkers are then correlated with the participants’ behavioral indications and used to classify experiences with or without active indication of behavior. However as no-report paradigms relied on conventional AFC methods, they too suffer the same limitations and have a number of other confounders e.g., pupil size changes used as no-report biomarker can be affected by different perceptual states regardless of perceptual alternation, or that eye-movements may be triggered due to piecemeal rather than exclusive percepts. (15,16)

Notice that all the above methods rely on two or more pre-defined categories for the participant to report by AFC and are based on the assumption that the categories defined by the experimenter are the same as those experienced by all participants. This assumption may be false, especially for atypical populations. Furthermore, the dynamics of transitions among states cannot be measured sensitively with button responses, which can only indicate abrupt transitions. Furthermore, these methods do not estimate an observer’s interpretation of the experimenter’s description of categorical boundaries, e.g. “exclusive left-tilt”, “piecemeal” etc. (see (17) for review of methods).

To address these shortcomings, we recently developed a continuous method called *Indicate-Follow-Replay Me: Binocular rivalry* (InFoRM: Rivalry) that can generate *a priori* personalized estimates of perceptual introspection and captures the dynamics of perceptual changes. The data can also be re-analyzed between and within perceptual categories that have been used in previous studies. The endogenous changes in perceptual experience reported with InFoRM are validated against exogenous changes via a physical replay of stimuli (Figure 1B). (17)

The InFoRM method allows us to address many questions that cannot be studied with current approaches. For example, rather than assuming each participant experiences 2 or more pre-specified categories, we can examine *a priori* how many distinct categories were reported for each participant and experimental condition. In the present study, we investigated three contrast conditions that are known to affect binocular rivalry: bilateral low, bilateral high, and low versus high contrasts, see more details here.(17) To determine the *a priori* categories first, data for each trial, participant, and contrast condition were analyzed using an unsupervised machine learning approach (k-means), and determined the clusters for a range of 1-10, 25, 50, 100, 1000 k-means (example Figure 2A). Then, we measured separation of the clusters using Silhouette analysis (Figure 2B). Next, we used a two-parameter fit to estimate the minimal number of clusters necessary to generate well-separated clusters (Figure 2C) and repeated the procedure for all participants and contrast conditions (Table 1). Finally, we repeated the analysis to validate the method against the physical replay data from Phase 4. As shown in Table 1, averaged across trials, participants, and contrast conditions, perceptual rivalry and physical replay generated 10 ±8 and 10 ±7 optimal clusters, respectively, which are well-separated (silhouette value 0.62 ±0.06 and 0.62 ±0.06).

**Figure 2:**
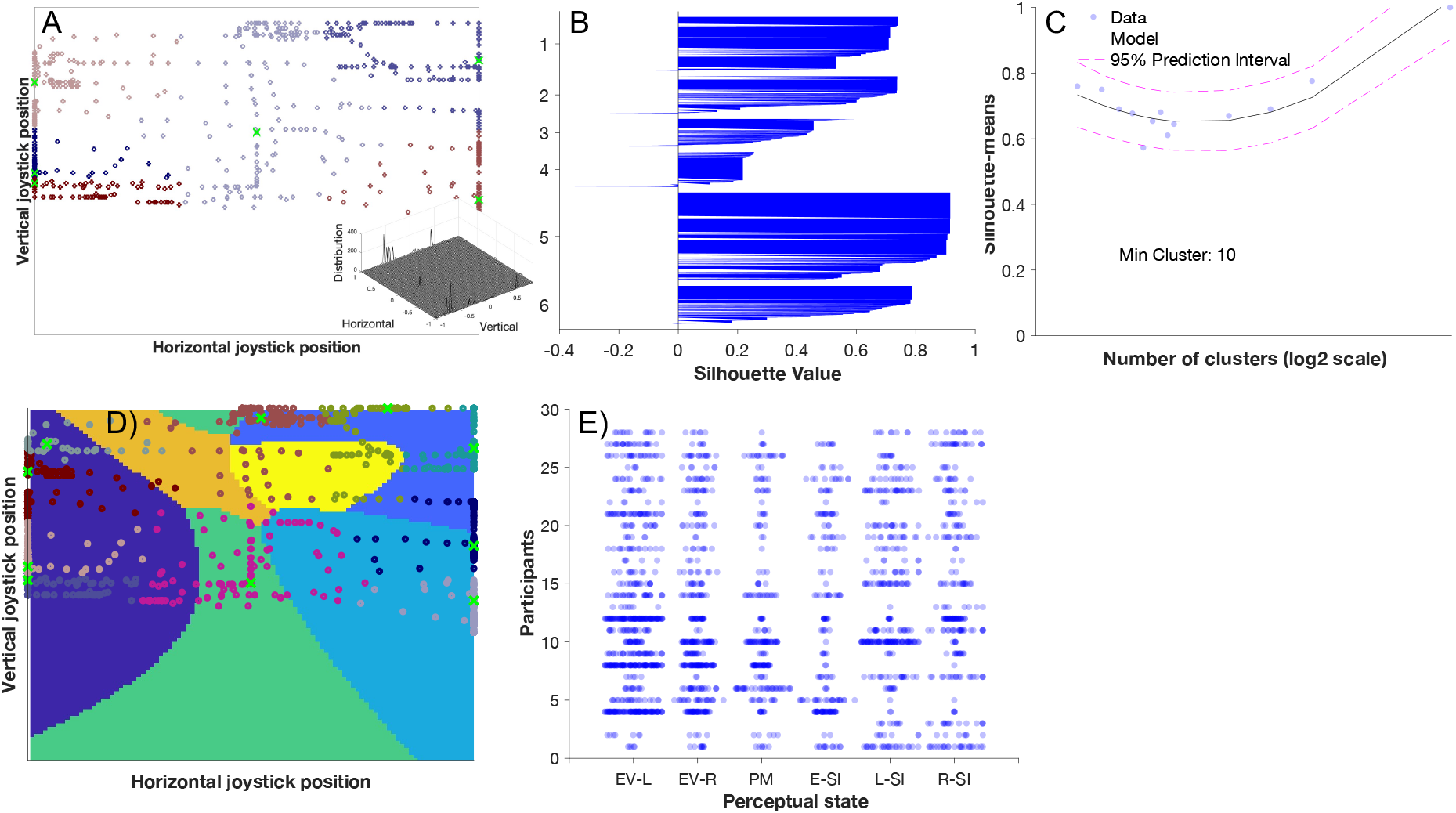
Data analysis using unsupervised cluster analysis. A) Example of joystick position during a Phase 3 rivalry trial. Depicted are raw data (dots), classified by k-means clustering illustrated by different colors. The centroid of each cluster is indicated via green x. B) The same data as in A) plotted with silhouette analysis, the separation of each data point is expressed with a silhouette value. C) Silhouette values were calculated for 1-10, 25, 50, 100, 1000 clusters. Then, the mean silhouette was calculated for each participant and cluster condition (blue dots) and fit with a second order polynomial (black line, magenta dashed lines show 95% confidence intervals). The minimum of the function identifies the minimum numbers of clusters, here 10 clusters. D) Illustration of raw data from A) with their optimal number of clusters (indicated with different hues) and centroids (green x) superimposed with that individual’s perceptual state map generated during phase 2 ‘Follow me’; (blue-left: exclusive left; green: equal superimposition; beige: superimposition with left-tilted predominance; blue-middle right: piecemeal; blue-upper right: exclusive right; yellow: superimposition with right-tilted predominance). E) Swarm plot of clusters for the low contrast condition for 8 trials for all 28 participants. Individual optimal k-means are superimposed on their perceptual state map, assigning number of k-means centroids for each of six perceptual states (x axis) for each individual (y axis).

**Table 1:** Summary of k-means optimal cluster analysis. Shown are mean and standard deviations across participants outcomes for the minimum silhouette values and their corresponding number of k-mean clusters for joystick report during perceptual and physical stimulus changes.

| Contrast condition | Perceptual Rivalry (Phase 3) |  | Physical Replay (Phase 4) |  |
| --- | --- | --- | --- | --- |
|  | Min. silhouette value | Min. k value | Min. silhouette value | Min. k value |
| Low vs. Low | 0.64 ± 0.055 | 9 ± 7 | 0.63 ± 0.065 | 10 ± 7 |
| High vs. High | 0.61 ± 0.054 | 12 ± 7 | 0.61 ± 0.059 | 11 ± 8 |
| Low vs. High | 0.62 ± 0.063 | 9 ± 8 | 0.64 ± 0.075 | 9 ± 7 |
| Mean | 0.62 ± 0.057 | 10 ± 8 | 0.62 ± 0.060 | 10 ± 7 |

Although individuals were trained on 6 categorical states (based on a review of previous studies), the results show that on average more distinct clusters experiences were perceived during rivalry. Our data allow us to examine the agreement between the six canonical states that are commonly assigned and the 9 or 10 clusters that participants spontaneously report. To answer this question, we return to the introspection maps that were created during InFoRM’s Phase 2 and superimposed these with the optimal k-means from Experiment 1 for each participant and condition. These maps were created based on each participant’s estimate of each of the six canonical categories previously described in the literature.(17) We assigned each k-means centroid from each trial to the closest of the six canonical categories and repeated this for each contrast condition (see example in Figure 2 D). As show in Figure 2E for the low contrast condition, the number of centroids in each perceptual state region varied between participants and occurred primarily in the *exclusive* portions of the joystick space as well as in the *superimposed* states with predominance of either left or right with fewer reports around *equal superimposition* or *piecemeal* observations during rivalry. These results suggest that *piecemeal* percepts can be thought of as an intermediate phase between both exclusive states (i.e. monocular sites) and superimposed states (binocular site).

Averaged across trials and participants, 13 ±9, 15 ±10, 12 ±11 centroids emerged for the low, high, and low vs. high contrast conditions, respectively, and were not significantly different from each other [repeated measure ANOVA, Greenhouse-Geisser *F*(2.0,53.1)=1.7, *p*>0.05, 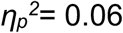]. However, as can be seen exemplarily in Figure 2E for the low contrast condition, the number of centroids across states varied significantly (two-way ANOVA, [*F*(3.0,80.0)=14.5, *p*<0.001,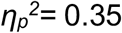]) but not was not affected by the contrast condition [*F*(2.0,53.1)=1.7, *p*>0.05, 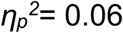].

We calculated the size of each classified area, as shown in Figure 2E, to investigate the relative sizes of different perceptual states. Averaged across trials, participants and contrast, the six states occupied 23% ±4, 16% ±5, 25% ±7, 9% ±7, 12% ±3, 15% ±3 and showed a significant difference. (17) The number of cluster centroids falling within the 6 classic categories, averaged across all levels, was 22 (exclusive left 27% of all clusters) ±25, 14 (exclusive right, 17%) ±15, 9 (piecemeal, 11%) ±13, 6 (equal, 7%) ±8, 19 (left-predominant superimposition, 24%) ±23, 10 (right-predominant superimposition, 14%) ±13. Interestingly, these results show that area and number of clusters mismatch for piecemeal and predominant left superimposed areas. In fact, although the piecemeal area of the introspection maps was the largest classification area overall, it housed only a small proportion of clusters. Taken together, a considerable number of clusters are generated in superimposed mixed states that resulted in 35% of all superimposed experiences and 12% piecemeal perception as reported previously. (17) These results suggest that current standard 2-3AFC methods have neglected these superimposed categories and thus may not accurately represent the experiences or their underlying neural site(s). Some studies have reported superimposition as a perceptual category during binocular rivalry, (18,19) but only a few used 4-5AFC methods to investigate the perceptual dynamics during binocular rivalry. (7,8) Only two studies have reported explicit experiences of superimposition with a predominance one eye’s stimuli. In one case it was invoked due to difference in spatial frequency (20) in the other it was invoked using the same spatial frequency but varying unilateral and bilateral stimulus contrasts. (17) Our results show that, even with bilateral equal stimuli, these experiences can emerge. It may be possible that studies that used ambiguous instructions such as ‘predominance’ may have captured instances of these experiences as well. (21,22) Importantly, while exclusive perception (global) and piecemeal (local) are thought to be a result of mutual inhibition of monocular sites, (23) superimposed percepts may activate distinct neural correlates (24) that might include binocular cells as suggested by different psychophysical investigations. (7,8,18)

In addition to investigating the categorical experiences during binocular rivalry, we next interrogate the transitory dynamics among these experiences during binocular rivalry.

First, the actual transition path for each trial (example in Figure 3A) was used to calculate the mean transition pathway (Figure 3B) for each participant and condition. Then, the most likely transition pathway for each participant and condition was estimated using a Hidden Markov Model (Figure 3C). Next, the similarity of the model with the actual data was estimated by using cross-correlation to determine the agreement between the model and data (Figure 3D). The resulting similarity measures (maximum correlation coefficient, lag at maximum correlation, and area under the curve) were further analyzed. Separate repeated measure ANOVAs were performed to test for an effect of contrast condition. An effect for area under the curve [*F*(1.8,48.0)=2.8, *p*<0.01, 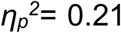] was found due to less AUC for the high vs low contrast condition. Maximum correlation coefficient [*F*(1.7,46.3)=1.4, *p*>0.05, 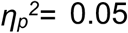], nor for lag at maximum correlation (bias) [*F*(1.0,27.0)=1.0, *p*>0.05, 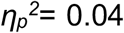] was found. The lag at maximum correlation was close to zero (1 ±0.11).

**Figure 3:**
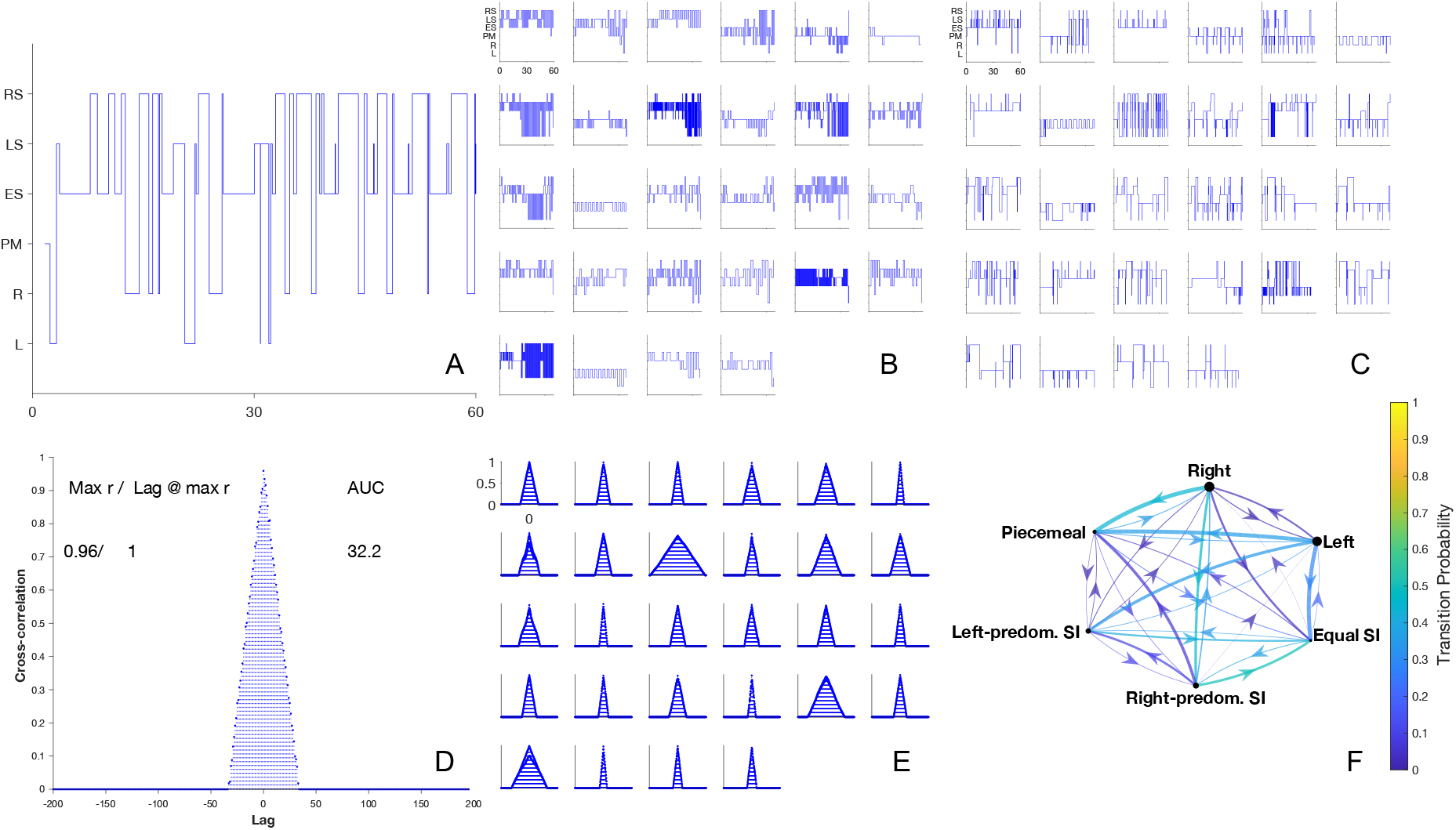
Analysis of binocular rivalry dynamics. A) A typical transition path of a single trial for one participant classified into 6 classic perceptual states (‘L’ left-tilted exclusive, ‘R’ right-tilted exclusive, ‘PM’ piecemeal, ‘ES’ equal superimposition, ‘LS’ left-tilted predominant superimposition, ‘RS’ left-tilted predominant superimposition) changes across time. B) Mean transition path for each participant during the low contrast conditions were then used to estimate the most likely transition path (C) calculated by a hidden Markov model. D) Cross-correlation for one participant as a function of lag between actual and model data. Maximum correlation coefficient, r, and its lag location relative to the optimum 0 lag as well as the estimated area under the curve (AUC) are included. E) Cross-correlations for each individual during the low contrast condition. F) Transition probability chain plot averaged across trials and participants for the low contrast condition.

Previously, we introduced a new way to analyze multistability data, which combines Markov chains and their ability to depict states, their connections, and the likelihood of each connection with the temporal priors, i.e., the mean duration of a percept before it transitioned to another state (indicated via arrow thickness, where arrow thickness increases with percept duration) and the mean duration of each principal state (nodes, where diameter increases with mean duration (3)). This method therefore makes predictions of state connections and their likelihoods and also incorporates temporal legacy of each of these transitions. The node diameters (Figure 3F) symbolize each state’s mean duration, each arrow thickness (weights) indicates the mean duration of a given perceptual state prior to transition to another state. We correlated the weights with the transition probability values for each contrast condition and found no correlation for the low (R: 0.01; p>0.05) or high contrast conditions (R: 0.13; p>0.05), but a positive correlation when using different dichoptic contrasts i.e., the longer the prior duration of percepts the greater the transition likelihood between these two perceptual categories (R: 0.39; p<0.05). On one hand, these results imply that when using equal bilateral contrasts, rivalry transition dynamics are not dependent upon prior accumulative experiences (weights), suggesting a primary role of intrinsic noise as driver for transition. (25) On the other hand, the positive correlation between weights and transition probabilities when using unequal bilateral contrasts suggest a role of prior experience, supporting the hypothesis that this type of multistable vision is explained by self-adaptation models. (26)

We compared probability distributions between contrast conditions using a Kullback Leibler divergence. As expected, when comparing the dissimilarity of transition probabilities for the equal bilateral contrast conditions (low/low vs. high/high), dissimilarity was the lowest (range: 0.05-0.29), whereas dissimilarity was higher for unequal bilateral contrast conditions (low vs. low/high conditions [0.11-0.50]; and high vs. low/high conditions [0.14-0.57]). As illustrated for the bilateral low contrast condition in Figure 3F, changes away from exclusive states to predominant-superimposed states are more likely and with longer mean durations (thicker arrows) compared to other changes. One hypothetical reason for this result could be the joystick arrangement i.e., left tilt for left exclusive and right tilt for right-exclusive percepts, however, as shown in Figure 3B and 3C, mixed states were not mere transit-states between two exclusive percepts for the majority of participants. As for data clustering, we show considerable individual differences in transition dynamics between perceptual states. Furthermore, the analysis reveals a higher transition probability between exclusive and left and right-predominance superimposed states. Specifically, for the low contrast condition the minimum transition probability 0 (no transition from left exclusive to right predominant superimposition, SI); maximum 0.50 (right predominant SI to equal SI), and a mean of 0.19. The results for high contrast [*min*: 0.007;(exclusive right to equal SI); *max*:0.50 (right-tilted SI to equal SI), *mean*: 0.18] and for the low versus high contrast conditions [*min*: 0.01(low contrast to high-contrast predominant SI); *max*: 0.50 (low contrast to low contrast predominant SI); *mean*: 0.19] indicate that transitions were more likely to occur between exclusive monocular and fused binocular percepts.

In conclusion, the combination of a continuous psychophysical approach, introspection estimates, and unsupervised cluster analysis revealed that on average more perceptual categories arise during binocular rivalry than previously thought. Moreover, binocular rivalry transitions are more likely to occur between exclusive and superimposed perceptual states than other state changes and are affected by prior experiences only when the interocular inputs are different. Together, these results suggest that conventional binocular rivalry paradigms do not capture the full range of experiences during binocular rivalry or their dynamics. Furthermore, transitions among states show greater variability than previously thought, in particular within superimposed perceptual categories. The results of the transition probability analysis imply that perceptual competition during binocular rivalry that is evoked by low-level stimuli arises as a conflict between monocular and binocular neural sites rather than mutually inhibiting monocular sites.

## Methods

The experiments were carried out in the facilities of Northeastern University, Boston. Written and verbal information about the project were provided in advance to the participants and they gave written informed consent before taking part. Ethics approval to conduct the experiments on human participants was in line with the ethical principles of the Helsinki declaration of 1975 and ethics board of the Northeastern University. The methods regarding the InFoRM Rivalry method have been reported elsewhere in detail. (17) Here we report methods and materials specific to the data analysis. Matlab (Mathworks, version 2023b) was used for data collection, analysis, and visualization of the results in the current study. Stimuli were presented on a LG 3D polarized monitor with a spatial resolution of 1920*1080 pixels in combination with radially-polarized LG cinema 3D glasses (AG-F310), 60Hz refresh rate and mean luminance of 61.9 cd/m^2^, and a Dell computer (Optiplex 7060). The viewing distance was 150cm. The participants wore radially-polarized LG cinema 3D glasses (AG-F310) and provided responses with a Logitech ExtremeTM 3D pro (Logitech Europe S.A.) joystick.

Binocular rivalry was induced in 28 normally-sighted participants using orthogonally oriented (±45°) sinusoidal gratings (2° Ø, 2c/°). Three contrast conditions (low versus low; high versus high, and high versus low) were tested in counterbalanced order. Raw data consisted of 3600 data points (60Hz joystick data sampling * 60seconds testing; 16.7ms temporal resolution) per trial (8 per contrast condition) that consisted of 2D joystick position estimates for each Phase 3 (rivalry) and Phase 4 (replay) and were stored in .*mat* files. Perceptual introspection maps and state assignment during Phase 3 (rivalry) and Phase 4 (replay) were described elsewhere. (17)

### Cluster Analysis

Horizontal and vertical joystick vectors were converted into Euclidean space for the Phase 3 (rivalry) data for each trial. Second, unsupervised clustering was performed for a range of clusters (1-10, 25, 50, 100, 1000) using the *kmeans* function applying the ‘cityblock’ method for each trial and averaging the results across trials for each condition. Third, we evaluated the separation among clusters using the *silhouette* function and applied again the ‘cityblock’ method. We found that the overall silhouette values were all positive, i.e. well-separated. As the choice of k-means is arbitrary, we decided to find the minimum separation value required, which represents the optimal clustering value. Hence, for the fourth step, we plotted the resulting silhouette values against k-means for each participant and for each condition, fit a quadratic function using *polyfit* and *polyval* functions to the data to estimate the minima of the fit, and extracted the corresponding optimal silhouette value and optimal number of k-means clusters. We repeated the above-described analysis for Phase 4 (replay).

Each participant’s optimal k-means value was used for the assignment to their respective introspection maps to find out where within the classification space the centroids would cluster. Then, we assigned each centroid for each trial with one of the six introspection classifications derived from previous binocular rivalry studies. For example, if a centroid arose in the introspection map area of ‘left exclusive”, that centroid was counted for left exclusive. This was repeated for each trial, participant, and contrast condition. SPSS software (IBM, version 28.0.0.0.(190)) was used to perform repeated measure ANOVAs.

### Transition Probability Analysis

#### Actual and HMM most likely transition path

Each trial’s perceptual state vector (3600 data) consisted of up to six distinct states and was averaged across trials for each participant to generate the average transition path. The *hmmestimate* function was used to calculate the mean transition probability for that trial. The *hmmestimate* function was repeated with the ‘pseudotransition’ setting using the mean transition value as some transition probabilities were very low. Next, the HMM most probable transition path for each trial was estimated using the *hmmviterbi* function. Single transition paths were visualized using the *stairs* function.

#### Cross correlation of Actual and HMM transition path

The *xcorr* function (‘normalized’ mode, maximum lag of ±200 time points = ±3.3 seconds) was used for the cross correlation of the actual mean paths and means of the most likely paths HMM paths for each participant, and contrast condition. The peak of the resulting cross correlation function, lag, and area under the curve (estimated using the *trapz* function) were taken.

#### Markov chains

The *hmmestimate* function was used to estimate the transition likelihoods between states for each trial, participant, and contrast condition. We used the *dtmc* function to estimate Markov chains that were then plotted using the *graphplot* function for each contrast condition.

As previously described, (3) we also included temporal legacy in the chain plot, indicated by increasing node diameter for mean durations and thicker arrows (weights) for longer prior mean durations before a transition occurred. Each weight was measured for each trial as a mean duration of how long either of the six canonical perceptual states lasted. The results were then averaged across trials and participants for each contrast condition. The *corrplot* function using Pearson’s method was applied for linear correlations between weights and transition probabilities, testing for R and for statistical significance test p. Kullback-Leibler similarity analysis was performed to compare the transition probabilities between contrast conditions applying the *KLDiv* function.

## Acknowledgment

Supported by NIH grants R01 EY029713 and R01 EY032162.

## Authors contributions

JS and PB developed the principal concept of the study and invented the InFoRM method. Both authors developed and refined the code of the four phases. Both authors contributed to the experimental design of the study. JS and a research assistant carried out the optometric screening and collected data. JS wrote the analysis code and the first draft of the manuscript. Both authors critically revised both code and the manuscript before submission.

## Competing interests

InFoRM was invented by PJB and JS and is disclosed as patent (pending) held by Northeastern University, Boston USA.

## Financial interests

Both authors are founders and shareholders of the company PerZeption Inc. (USA) which has licensed the patent for InFoRM.

## Data availability

Data generated for this study and code can be found here: **10.5281/zenodo.13831435**.

## Notes

https://zenodo.org/records/13831435

